# Collective gradient sensing by dilute swimming bacteria without clustering

**DOI:** 10.1101/2022.11.17.516991

**Authors:** Tatsuro Kai, Takahiro Abe, Natsuhiko Yoshinaga, Shuichi Nakamura, Seishi Kudo, Shoichi Toyabe

## Abstract

We characterize the taxis enhancement of swimming bacteria by collective migration without apparent clustering. We confine dilute *Salmonella* suspension in a shallow channel and evaluate the thermotaxis response to local heating and diffusion. By combining cell tracking analysis and numerical simulation based on simple modeling, we show that the alignment interaction suppresses orientation fluctuation, strengthens migration bias, and also prevents the dispersion of accumulated population. The results show a prominent example of how a collective motion of active matter implements a biological function.

---

Nearly all bacterial species implement the taxis ability to migrate along the gradient of, for example, chemicals and temperature to seek a better environment. The mechanism underlying taxis differs from species to species. Bacteria with peripheral flagella such as *Escherichia coli* and *Salmonella* exhibit biased random walk with alternative run and tumble motions [1] [Fig. 1(a)]. Signal-dependent modulation of the tumbling frequency enables taxis. However, since such trial-and-error swimming of individual cells is highly stochastic, the taxis efficacy is not expected to be high. On the other hand, it is known that bacteria suspended at high density swim collectively and exhibit rich spatiotemporal phenomena such as long-distance ordering [2, 3], giant number fluctuations [2], and vortex formation [4, 5] caused by interaction among cells. A question that naturally arises is how the cellular interaction affects the taxis performance.

**FIG. 1.**
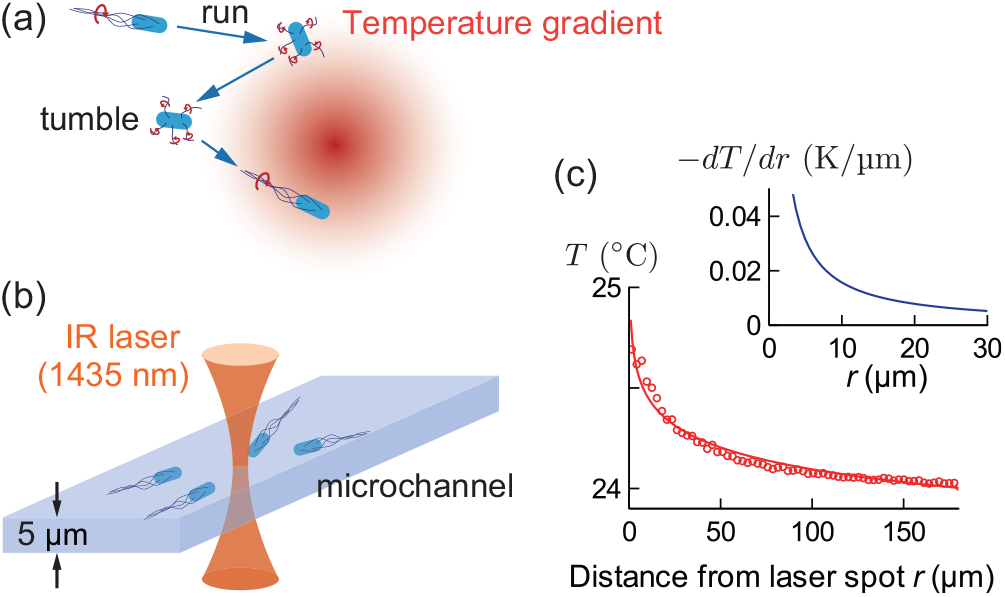
Bacterial taxis. (a) Stochastic run and tumble motion realizes taxis. (b) Experimental setup. An infrared (IR) laser was focused under a microscope to generate a temperature gradient in a microchannel [Fig. S1 ]. (c) Temperature gradient generated by local laser heating with 200 mW power. The profile was well fitted by *a*− *b* ln *r* with *a* = 24.8 and *b* = 0.157. *r* is the distance from the laser spot. The inset is the gradient calculated as *b/r*. See Fig. S4 for details.

It has been observed that taxis is enhanced with increasing cellular density in a variety of systems including dense bacteria swarming on agar plate [6] and migrating eukaryotic cells [7, 8]. The density dependence suggests that the cellular interaction affects the taxis behavior. In such systems, cluster formation is supposed to be the primary mechanism. A cluster, which effectively behaves as a large body, makes it easier to sense a gradient or even enables gradient sensing even if individuals can sense only a concentration instead of a gradient.

In contrast, it was demonstrated that the chemotaxis performance of dilute swimming bacteria increases with the cell density at a low cell density where cluster formation is not expected [9], while a further increase in density suppresses the taxis due to collective motion. There might be a novel enhancement mechanism for dilute bacterial suspensions, where the exclusion volume effect does not dominate the interaction, but it remains elusive. This work aims to quantitatively characterize how cellular interaction modifies taxis at low cell density and clarify its mechanism. For this aim, we focus on a low-density cell suspension (1∼ cell/100µm^2^) of *Salmonella enterica* cells, where apparent clustering is not observed. As well we used thermotaxis, that is, the property that *Salmonella* migrate along a temperature gradient. Dynamic and local heating by laser irradiation allows us to measure the temporal dynamics of the taxis response instead of the steady behavior. This helps us to understand the detailed taxis enhancement mechanism by cellular interaction.

In a dilute system, the taxis may be evaluated based on a framework similar to that for the thermophoresis of non-active microscopic objects [10]. Let 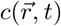 be the particle density at position 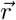 and time *t*. Under a given temperature gradient 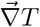, both the taxis and the thermophoretic flow of non-active particles are phenomenologically described as

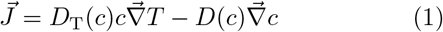

in a linear-response regime. *D*_T_ and *D* are thermodiffusion and diffusion coefficients, respectively. The balance between migration bias and diffusion quantified by the Soret coefficient 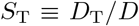 determines the efficacy of detecting the temperature gradient. Actually, if the dependence of *D*_T_ and *D* on *c* are negligible, we obtain the steady distribution 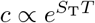 by solving 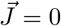. This is often the case for thermophoresis. The phoresis and taxis have distinct origins. The thermophoresis is basically driven by the interaction between the particle surface and solvent [11], whereas bacteria realize thermotaxis by modulating their swimming pattern based on biochemical responses to a thermal gradient[1]. Bacteria swim ballistically at a short time scale and effectively exhibit diffusional motion only at a long time scale[1]. Despite these differences, it would still be effective to evaluate the taxis efficacy based on *D*_T_(*c*), the effective diffusion coefficient *D*^eff^ (*c*), and their ratio 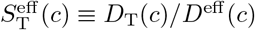

We observed the swimming cells confined in a thin microchannel with the height of *h* = 5 µm, which is comparable to the cell dimensions (2–3µm in length and ∼ 1 µm in width) [Fig. 1(b)]. Cells may pass each other. However, since vertical swimming is negligible, we treat the system as two-dimensional and evaluate *c, D*_T_, and *D*^eff^ as two-dimensional quantities. Irradiation of focused infrared laser under a microscope generated an axisymmetric pseudo-two-dimensional temperature gradient [12, 13]. This setup enables the instantaneous switching on and off of the gradient for probing the taxis dynamics. The thermal convection due to the local heating was negligible because of the reduced channel height. The consumption of nutrients by cells creates a nutrient gradient and makes the dynamics complex [13, 14]. We used the observation buffer lacking nutrients to focus on the effect of the cellular interaction on taxis. We also evaluated *D*_T_ using the initial migration speed since cellular respiration may form the spatial oxygen gradient at a later time. The magnitude of interaction was controlled by varying the cell density.

## Results – taxis

The spatial temperature profile under local laser heating was well fitted by a logarithmic function as expected from the two-dimensional open system with a single heat source[15] [Fig. 1(c)]. The maximum temperature increase was about 0.7 K.

The cells migrated along the temperature gradient 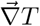 and accumulated in the vicinity of the heating spot [Figs. 2(a), S7], indicating *D*_T_ > 0 under the present condition. The cells dispersed when the laser was turned off. The mutant strains without flagella (fla-) nor Che system (che-[16]) did not accumulate significantly [Fig. S8]. The che- strain produces an intact flagellar apparatus but rotates its motors exclusively counter-clockwise without tumbling. Hence, the migration originates in the taxis, not in the optical-tweezers effect [17] nor the thermophoresis [10].

**FIG. 2.**
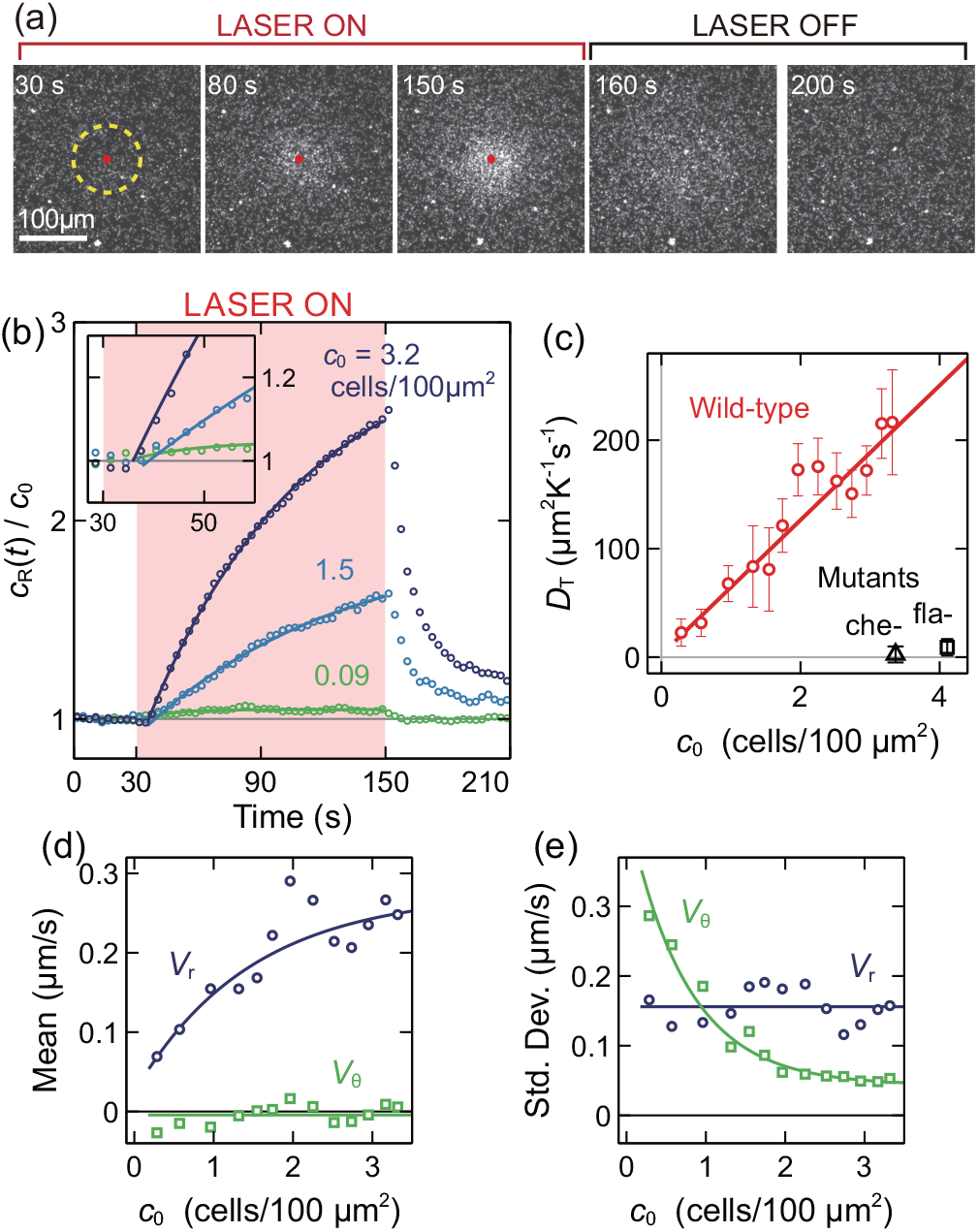
Taxis to local heating. (a) The snapshots of the migration at *c*_0_ = 2.5 cells*/*100µm^2^ were observed by darkfield microscopy. The red dot indicates the laser irradiation location. (b) The normalized cell density *c*_*R*_(*t*)*/c*_0_ in the central region within *R* = 50 µm (yellow dotted circle) in (a). *c*_*R*_(*t*)*/c*_0_ was evaluated as the ratio of the light intensity scattered by the cells under dark-field microscopy. *c*_0_ was separately evaluated by identifying individual swimming cells. The data are fitted by exponential curves (solid lines) in the range between 40 s and 150 s. The time profiles of 20 runs with similar *c*_0_ are averaged. (c) Dependence of *D*_T_ on *c*_0_. The solid curve is a linear fitting. 141 independent observations (wild-type) are sorted by *c*_0_, divided into groups containing 20 data with 10 overlaps, and analyzed in each group. *N* = 7 for each mutant. The error bars indicate the standard errors. (d, e) Single-cell tracking. The mean (d) and standard deviation (e) of radial (*V*_r_) and angular (*V*_*θ*_) components of the velocity under the local heating averaged in the distance range between 74 and 198 µm from the heating spot and the time range of between 40 and 70 s. See Fig. S9 for the mean swimming velocity and SI Method for details. Each point is obtained from 20 runs. The solid curves are exponential or constant fittings. The plot was limited to *c*_0_ *>* 0.1 cell*/*100µm^2^ due to limited sample number at low *c*_0_.

Let *c*_*R*_(*t*) be the mean cell density in the vicinity of the heating spot (the region within *R* = 50 µm) and *c*_0_ be the mean cell density in the same region before the laser is turned on. *c*_0_ was typically in the order of 1 cell/100µm^2^. The normalized cell density *c*_*R*_(*t*)/*c*_0_ increased linearly with a short time delay and saturated with the timescale of ∼ 80 s [Fig. 2(b)]. The short delay was supposedly caused by the response delay of individual cells and the large *R* we used.

The increase in *c*_*R*_ should equal the influx through the region’s boundary. The magnitude of the influx is obtained by integrating the radial component *J*_*r*_ of 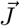 at the peripheral, *ċ*_*R*_ = 2*πRJ*_*r*_/*πR*^2^. Initially,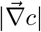 is negligible because of the uniform cell density. Hence, Eq. (1) is reduced to 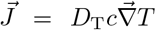, which leads 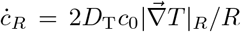. Here,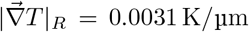 is the temperature gradient at *R*. Thus, we can evaluate *D*_T_ as *D*_T_ = *αċ*_*R*_(*t*_0_)/*c*_0_ with 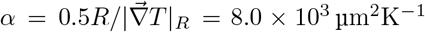. Here, *t*_0_ = 30 s is the time that the laser irradiation starts. We fitted the time profile of *c*_*R*_(*t*) during heating by an exponential curve and evaluated *ċ*_*R*_(*t*_0_). We excluded the first 10 s during heating from the fitting range to remove the effect of the short time delay of *c*_*R*_(*t*) rising. We found that *D*_T_ increases with *c*_0_. Since the increase in *c*_0_ increases the contact frequency of cells, the results imply that the cellular interaction enhances taxis [Fig. 2(c)].

For further characterization of migration dynamics, we tracked individual cells and evaluated the radial (*V*_r_) and angular (*V*_*θ*_) components of the swimming velocity under the temperature gradient [Fig. 2(d, e)]. Since it was not feasible to track all the cells due to their overlaps, the analysis was limited to a subset of the entire population. The mean of *V*_r_ increases with *c*_0_ while the standard deviations of *V*_*θ*_ decrease with *c*_0_. In contrast, the mean swimming velocity does not significantly change with *c*_0_ [Fig. S9 ]. These results imply that the cellular interaction strengthened the directional bias toward the heating spot by aligning the swimming direction without affecting the swimming speed. Reliable evaluation of *D*_T_, *V*_*r*_, and *V*_*θ*_ at vanishing *c*_0_, which should converge to the taxis of a single cell, is challenging and was not possible in the present experiments due to poor statistics.

## Results – cellular interaction

The above results indicate that cellular interaction enhances the taxis. To characterize the interaction, we analyzed the swimming trajectories used in Fig. 2(d,e) and evaluated the collision angles *θ*_in_, *θ*_out_, and their difference Δ*θ* = *θ*_out_−*θ*_in_ in the absence of thermal gradient [Fig. 3] [3, 18–20].

**FIG. 3.**
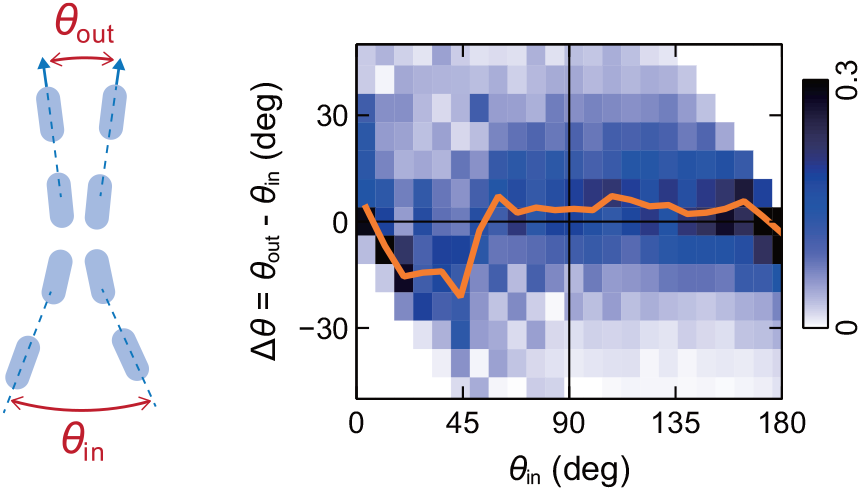
Alignment interaction quantified by the angular displacement by collision. We used the same data as that in Fig. 4 (*h* = 5 µm and *c*_0_ *<* 1.5 cells*/*100µm^2^). We analyzed 11,823 events where two cells come to close in a distance less than 2.2 µm. See SI for the details. The color intensity indicates the frequency of the collision events normalized in each column. The solid line indicates the peak locations of the skewed normal distributions fitted in each column.

The cells aligned after collisions with *θ*_in_ <45° and did not significantly align otherwise. Such mixed asymmetric interaction of polar and nematic types has been observed for the binary collision of the objects that can pass each other [20].

## Results – diffusion

Next, we evaluated how the modulation of diffusion by cellular interaction affects the taxis. After the laser was turned off, the cells dispersed quickly in an exponential manner [Fig. 2b]. The time constant of the dispersion *τ*_dis_ increased with *c*_*R*_ at the timing of laser off [Fig. S10]. We expect *τ*_dis_ to be independent of the density for passive Brownian particles unless the particles are extremely dense. The result implies that the cellular interaction modulates diffusion. However, the tendency of the dispersion rate to decrease with *c*_0_ is opposite to the taxis enhancement *c*_0_ seen in Fig. 2, where cells move in a similar direction toward the heating spot.

For more characterization, we tracked the diffusion of individual cells without laser heating. We mixed cells expressing green fluorescent protein (GFP) to track a long trajectory at high cell density. We first measured *c*_0_ by counting cells in phase-contrast observation and then quantified the diffusion using the GFP cells in the same observation area in succession.

The run and tumble motion generated ballistic and diffusive motions at short and long time scales, respectively [Fig. 4(a, b)]. Trajectories became more diffusive at higher *c*_0_, implying that the cellular interaction modulated the swimming pattern.

**FIG. 4.**
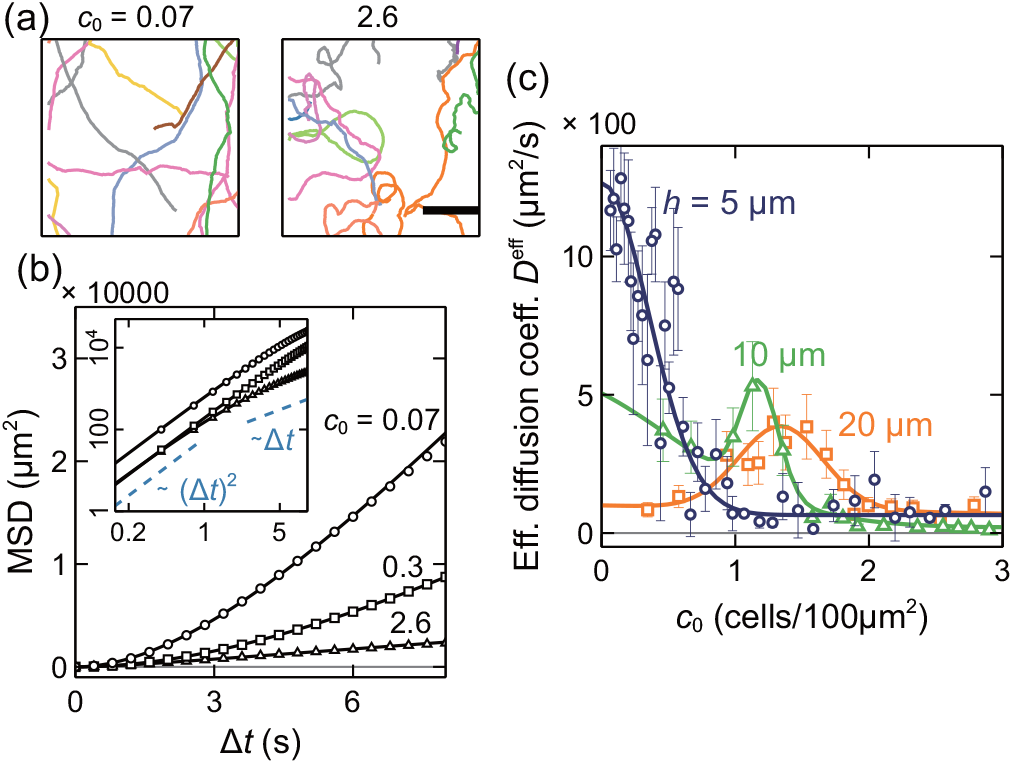
Diffusion in the absence of temperature gradient. (a) Two-dimensional trajectories in the chamber with the height of *h* = 5 µm. The scale bar indicates 100 µm. (b) The mean square displacement (MSD) curve in the chamber with the height of *h* = 5 µm. The values of *c*_0_ are indicated in the unit of cells*/*100µm^2^. The inset is the log-log plot. See Fig. S12 for the plots with error bars. (c) Effective diffusion coefficient *D*^eff^. The solid curves are fitting curves (see section VI of SI). The numbers of independent observations are 389, 120, and 192 for *h* = 5, 10, and 20 µm, respectively. The runs sorted by *c*_0_ are divided into groups of 10 (*h* = 5 µm), 8 (10 µm), or 12 (20 µm) runs and analyzed collectively in each group. The error bars indicate the standard errors.

The mean square displacement (MSD), 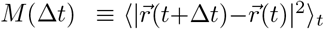, characterizes the diffusion [Fig. 4(b)]. Here, the average⟨·⟩ _*t*_ is taken for different *t*. The effective diffusion coefficient *D*^eff^ is defined in the long Δ*t* limit as *D*^eff^ = lim_Δ*t*→∞_ *M*(Δ*t*)/4Δ*t* in two dimensional systems. Since trajectories accessible by experiments are relatively short, we exploited a model fitting for estimating *D*^eff^. The MSD of a self-propelled particle with random directional change is described by

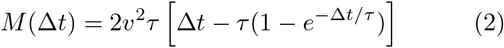

for a broad range of systems [21–23] including a run- and-tumble motion. Here, *v*is the swimming velocity, and *τ* is the mean duration between successive direction changes. In the present system, the change in direction is supposed to be caused by the tumbling and the cellular interaction. At large Δ*t*, Δ*t, M*(Δ*t*)≃2*v*^2^*τ*Δ*t*, which yields *D*^eff^ = *v*^2^*τ*/2. We fitted *M*(Δ*t*) by Eq. (2) to obtain *D*^eff^.

*D*^eff^ decreased with *c*_0_ as expected from the increase in diffusive motions. *τ* significantly decreased with *c*_0_, whereas clear dependence of *v*on *c*_0_ was not observed [Fig. S11]. Thus, the cells change directions more frequently at higher *c*_0_. The result is consistent with the extended dispersion time after the laser is off at high cell density [Fig. S10]. Once accumulated, the cells disperse slowly due to the slow diffusion caused by cell-cell interaction. These results indicate that the taxis enhancement by cell interaction is caused by both the increased migration bias and suppressed diffusion at the accumulated region.

We also measured *D*^eff^ in thicker channels with *h* = 10 and 20 µm as the references. At small *c*_0_, *D*^eff^ decreased with *h*. It is possible that the confinement to a shallow channel suppresses the directional change and enhances the speed. Interestingly, we observed nonmonotonic dependence of *D*^eff^ on *c*_0_ in the 10- and 20-µm channels [Fig. 4(c)]. Such peak was previously observed, but its mechanism has remained unclear [24].

This characteristic may be explained as follows. The alignment interaction suppresses the directional fluctuations and enhances the ballistic motion, which increases *D*^eff^. However, at high *c*_0_, due to the frequent collisions among cells and long-range hydrodynamic interaction, the correlated swimming may be destabilized. The balance between the alignment and destabilization may shape the peak of *D*^eff^ (*c*_0_). In the thick channels, cells can easily pass each other, and the destabilization would not be effective unless *c*_0_ is very large. Therefore, the peak position may shift to a large *c*_0_ region. Note that, due to the limited focal depth, *c*_0_ in 10- and 20-µm channels may be underestimated. However, this does not qualitatively affect the conclusion.

## Results – Soret coefficient

The effective Soret coefficient 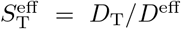 increased steeply when *c*_0_ exceeds a threshold cell density 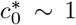 cell/100µm^2^ [Fig. 5]. The mean free path at 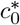 is roughly estimated a 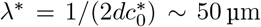, where *d* = 1 µm is the typical cell width. *λ*^*^ is comparable to the size of the heating region [Fig. 1(c)]. This is reasonable because the collective taxis enhancement is not expected if cells do not interact during the migration toward the hot region. The actual value of *λ*^*^ should be larger because cells can pass each other even in the present thin chamber. However, *λ*^*^ would still be comparable to the heating-spot size.

**FIG. 5.**
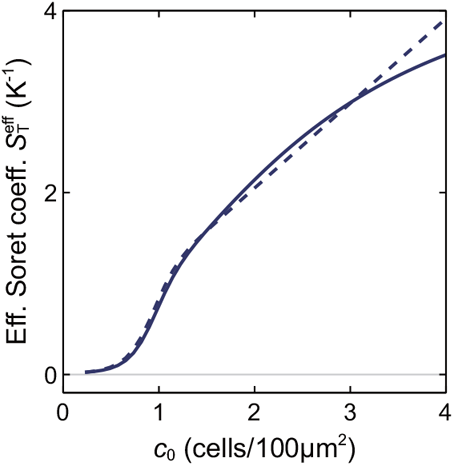
Effective Soret coefficient 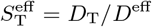 in the 5-µm chamber calculated based on the fitting curves of Figs. 2(c) and 4(c).

## Results – simulation

For elucidating the mechanism of the taxis enhancement, we modeled the cells as self-propelled spherical particles and performed numerical simulations [Fig. 6] [25–29]. See SI (Section VII) for details. The particles translate with a constant velocity and change their swimming direction by the interaction with other particles as well as rotational fluctuation with white Gaussian statistics. The interaction includes the short-range repulsion, which models the excluded volume effect, the polar interaction, which aligns the swimming direction of nearby cells, and the long-range hydrodynamic interaction, in which the cells are modeled as pushers. We also include the driving by an external field for measuring response.

**FIG. 6.**
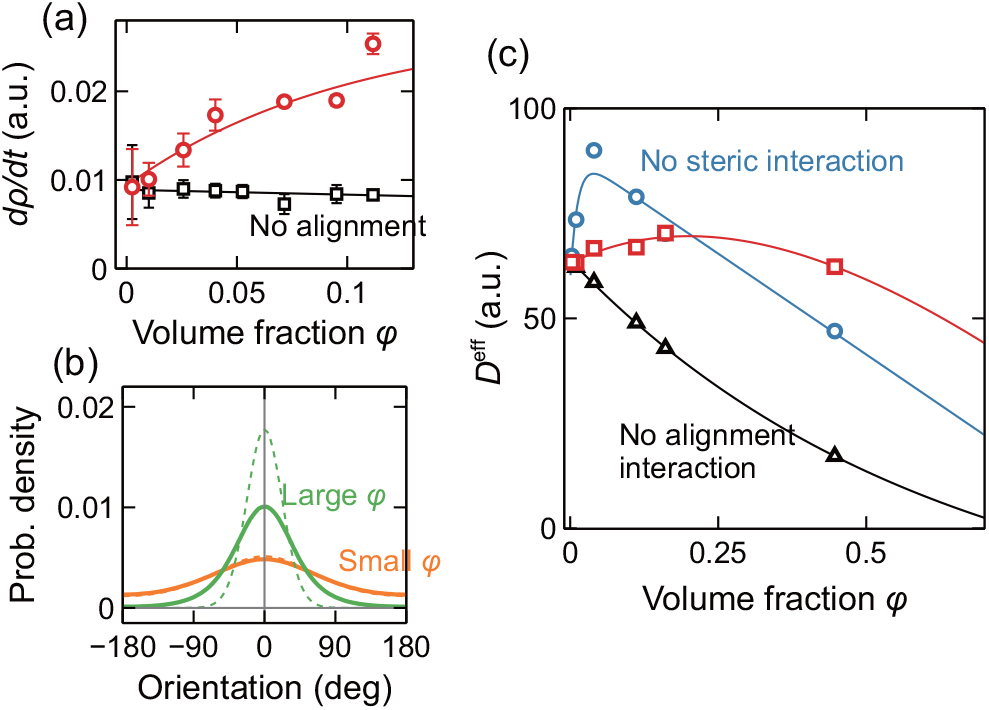
Simulation. (a) Response to a Gaussian-shaped axisymmetric external field. *dρ/dt* is the rate of increase of the density *ρ* within the region around the center (see SI for details). The solid curves are exponential fitting. (b) Steady distribution of velocity orientation obtained by simulations (solid lines) and mean-field approximation (dashed lines, see (S11)). Uniform external field is induced in the direction of *θ* = 0. *φ* = 0.11 (green) or 0.0025 (orange). See SI (Section VIIb) for the definition of response to an external field. (c) Effective diffusion coefficient with/without steric repulsion and polar interaction. The solid curves are exponential+linear (blue and red) or exponential (black) fitting. See also Fig. S6 for the effective Soret coefficient.

The simulations qualitatively reproduced the taxis and diffusion observed in the experiments despite the simplicity of the model [Fig. 6]. Specifically, the response to a Gaussian-shaped external field increased with the initial particle density measured by the volume fraction *φ* [Fig. 6(a)]. Such density dependence was not observed without alignment interaction. When a uniform external field was imposed, the steady distribution of the orientation became steeper with *φ* [Fig. 6(b)]. This result reproduced the experimental results that the mean of the drift velocity increased and its variance decreased [Fig.2(d, e)], validating the hypothesis that the alignment interaction enhances the migration bias and suppresses the rotational diffusion.

The peak of *D*^eff^ observed in experiments [Fig. 4(c)] was also reproduced by simulation [Fig. 6(c)]. Without the alignment interaction, *D*^eff^ monotonically decreased, validating that the cellular alignment enhances the diffusion. The peak of *D*^eff^ was large without the excluded volume effect, where the particles can slip through others. This situation models the experiments with thick chambers, in which we observed prominent peaks [Fig. 4(c)]. Let the volume of a single cell be *V* = 3 µm^3^. A typical cell density in the experiment *c*_0_ = 1 cell/100µm^2^ is converted to the volume fraction of *φ* = 0.006 for *h* = 5 µm channel. However, since the interaction range rather than particle size affects the cellular interaction more, it is not straightforward to compare *c*_0_ and *φ*.

## Discussion

Bacterial thermotaxis has been investigated since 1970s [30–32]. The density dependence of the thermotaxis was recently studied with a focus on the effect of the cellular metabolism [13, 14, 33, 34]. Cells’ secretion and consumption of chemicals such as glycine and oxygen create a chemical gradient, which induces chemotaxis and modulates the migration. Swimming speed variation by temperature also induces a temperature-dependent diffusion and enables a thermotaxis without receptors for taxis [34].

Here, we circumvented such complications and focused on how the cellular physical interaction affects the taxis behavior. As mentioned, the collective gradient sensing of dense cells has been explained by cluster formation [6–8]. In contrast, in the present dilute system, distinct cluster formation was not observed, implying a new class of collective sensing mechanisms. Based on the experimental results, the enhanced migration bias we demonstrated here may be explained by the following mechanism. The taxis of a single cell is not effective due to strong fluctuations. However, in a bacterial suspension, cellular interaction at repetitive instantaneous contacts aligns the swimming directions of the nearby cells to their major direction and increases the sensing limit.

The taxis at vanishing cell densities, corresponding to single cell taxis, was not well resolved due to low statistics. The simulation qualitatively reproduced the enhancement of taxis by cellular interaction. Quantitatively, however, the simulation [Fig. 6a] shows limited taxis enhancement compared to the experiment [Fig. 2c]. There may be other effects not considered in the simulation. For example, there may be a static interaction between cells that enhances accumulation. We have focused here on the essence of how the alignment interaction affects the taxis. More detailed modeling is left for future studies.

We thank Yusuke T. Maeda, Tetsuya Hiraiwa, and Yohei Nakayama for helpful discussions and Mao Fukuyama and Akihide Hibara for technical assistance. This research was supported by JSPS KAKENHI (JP18H05427 to ST and JP20K03874 and JP24K00591 to NY), JST ERATO Grant Number JPMJER2302, Japan, and Tohoku University Nanotechnology Platform sponsored by MEXT, Japan (JPMX09F-20-TU-0083).

