## Supplementary methods and figures. for "Collective gradient sensing by dilute swimming bacteria without clustering"

### Supplemental Information for
### "Collective gradient sensing by dilute swimming bacteria without clustering"

Tatsuro Kai,[1,*] Takahiro Abe,[1,*] Natsuhiko Yoshinaga,[2,3,4] Shuichi Nakamura,[1] Seishi Kudo,[1] and Shoichi Toyabe[1,†]

[1]*Department of Applied Physics, Graduate School of Engineering, Tohoku University, 980-8579 Sendai, Japan*
[2]*WPI Advanced Institute for Materials Research, Tohoku University, 980-8577 Sendai, Japan.*
[3]*MathAM-OIL, AIST, 980-8577 Sendai, Japan.*
[4]*Department of Complex and Intelligent Systems, Future University Hakodate, 116-2 Hakodate, Japan.*

#### I. CELL PREPARATION

The wild-type (SJW1103), GFP mutant expressing green fluorescent protein, che- mutant (SJW3076, Δ(cheA-cheZ)) lacking the che system[1], and fla- mutant lacking flagellar filament (SJW2936)[2] of *Salmonella enterica* were cultured in L-Broth medium for overnight at 37 °C, transferred to fresh L-Broth medium, and incubated for 5 (wild-type) or 8 hours (GFP mutant). The buffer was exchanged to an observation buffer (10 mM $KH_2PO_4$, 3 M KOH, 0.1 mM EDTA, and 10 mM sodium L-lactate) by centrifugation.

#### II. CHAMBER PREPARATION

The observation chamber was prepared as follows. Polydimethylsiloxane (PDMS, Sigma-Aldrich) with addition of 0.375 % DOWSIL 501W additive (DOW) that provides hydrophilic surface[3] was solidified on a mold built on silicon wafer for the 5-µm chamber. The PDMS was ripped from the mold, treated by plasma cleaning, and fixed on a coverslip surface. For the 10- and 20-µm chambers, we used both-adhesive tape (Teraoka) as the spacer between flat PDMS+501W ceiling and coverslip floor.

We flowed 10 mg/ml bovine serum albumin in observation buffer into the chamber for reducing the cell-surface interaction. Then, the bacteria suspension was flowed into the chamber.

#### III. OBSERVATION

The cell motions were observed under an inverted microscope (Olympus) equipped with 20× objective lens and CMOS camera (Basler) at 30 Hz or 50 Hz at the room temperature (25.0 ± 0.5 °C). The video was analyzed by a software developed in the lab on LabView (National Instruments). For the taxis measurement, laser with an wavelength of 1435 nm was irradiated through the objective lens for the local heating of the buffer.

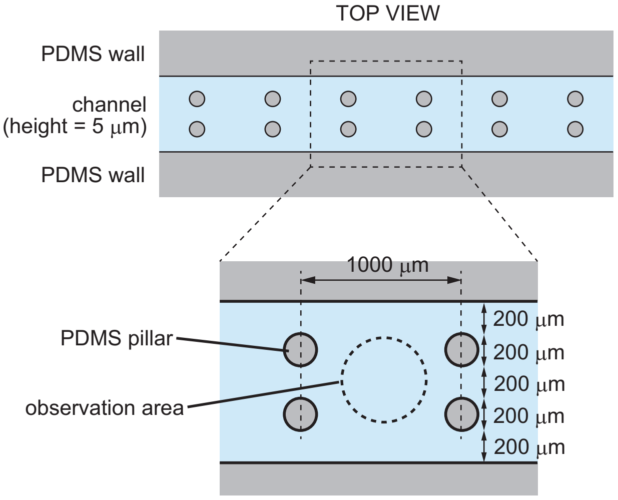


FIG. S1: Design of observation chamber.

#### IV. TEMPERATURE PROFILE

The temperature profile under laser heating was measured using two fluorescent dyes, Rhodamine B (Sigma-Aldrich) and Rhodamine 101 (Sigma-Aldrich). These dyes have fluorescence with different dependence on the temperature. The spatial temperature gradient may induce a thermophoretic migration of the dyes. However, the ratio of the two dyes provides the temperature assuming that the thermophoretic magnitude is the same for these two dyes. The calibration curve was measured using a real-time thermal cycler (BioRad) (Fig. S2). The $h = 5$ µm channel was first filled with 30 µg/ml Rhodamine B (Sigma-Aldrich) or 30 µg/ml Rhodamine 101 (Sigma-Aldrich) were flowed into the chamber. The spatial profile of the fluorescence of these solutions were measured with/without laser irradiation with different powers and analyzed to estimate the temperature profile (Fig. S3).

The profile was well fitted by $T(r) = a - b \ln r$, where $r$ is the distance from the laser spot. This function is the steady solution of the thermal diffusion equation in open two-dimensional system with a central heat source.

---

*These authors contributed equally.
†Electronic address:

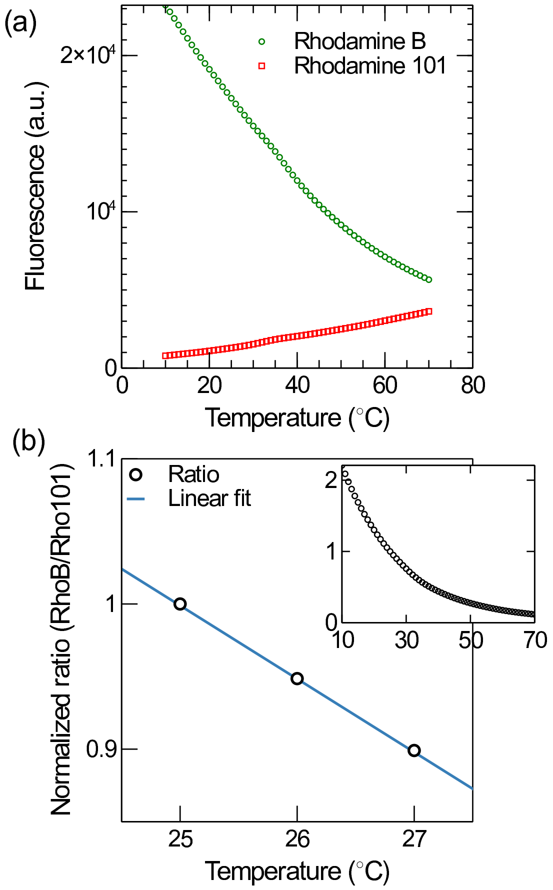


FIG. S2: Calibration curve for the temperature measurement. (a) The fluorescence of 30 µg/ml Rhodamine B (circle) and 30 µg/ml Rhodamine 101 (square). (b) The fluorescence ratio of Rhodamine B and Rhodamine 101 normalized by the ratio at 25 °C. The ratio was fitted in the range of 25 °C $\leq T \leq$ 27 °C by a linear curve $-0.505T + 2.261$.

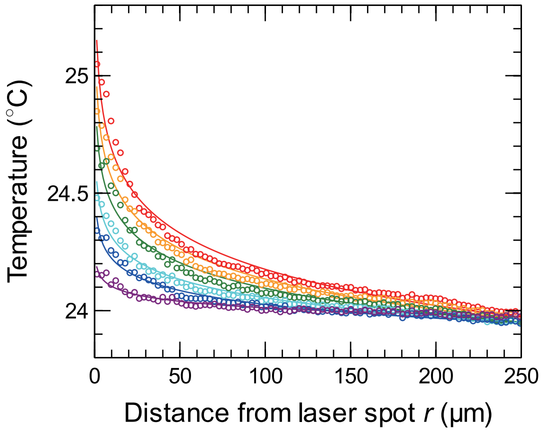


FIG. S3: Temperature profile under local heating with different laser power: 50 (purple), 100 (blue), 150 (cyan), 200 (green), 250 (orange), and 300 mW (red). The laser power indicates the power of the laser diode and larger than the actual radiation power to the chamber. The fitting curves are $T(r) = a - b \ln r$ with $(a, b)$ = (24.2, 0.0397), (24.4, 0.0849), (24.6, 0.113), (24.8, 0.157), (25.0, 0.187), and (25.2, 0.222) for 50, 100, 150, 200, 250, and 300 mW laser power, respectively.

#### V. CELL TRACKING

After particles are identified by image analysis, we recovered their trajectories by connecting particle pairs in successive two video frames. For each particle in $i$-th frame, we calculate scores for all particles within a threshold distance in the $(i+1)$-th frame and choose the particle with the least score in the $(i+1)$-th frame as the identical particle as the one in the $i$-th frame. If there is no particle with a score less than a threshold value, we stop the tracing of this particle. On the other hand, if a particle in $(i+1)$-th frame has no correspondence in the $i$-th frame, we start a new trace for this particle. The score is defined as the weighted summation of three components; (i) squared distance from the location predicted by optical flow method, (ii) squared distance from the location predicted by the particle velocity, and (iii) the area difference of the particle image. They have the weight of 1, 5, and 5, respectively.

We split the traces into 10-frame short traces. Let $\vec{r}_0$, $\vec{r}_1$, $\vec{r}_{\mathrm{m}}$, and $\vec{r}_{\mathrm{laser}}$ be the start, end, and mean locations of the short traces and the laser spot location, respectively (Fig. S4). The swimming speed is calculated as the magnitude of $\vec{v} = (\vec{r}_1 - \vec{r}_0)/9\Delta t$, where $\Delta t$ is the frame period. $V_r$ and $V_\theta$ are calculated as the components of $\vec{v}$ in the $\theta_{\mathrm{laser}}$ direction and $\theta_{\mathrm{laser}} + \pi/2$ direction, respectively. Here, $\theta_{\mathrm{laser}} = \mathrm{ang}\,(\vec{r}_{\mathrm{laser}} - \vec{r}_{\mathrm{m}})$ is the direction to the laser spot, where $\mathrm{ang}\,\vec{r}$ denotes the angle of a vector $\vec{r}$. The mean was taken for the cells in a window of time and the distance from the laser spot as indicated in the caption of Fig. 3.

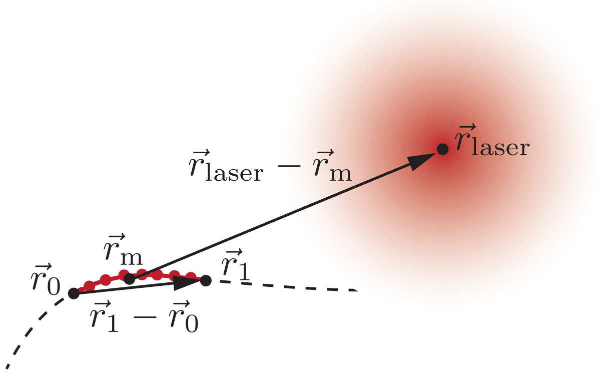


FIG. S4: Definition of the vectors. Red line is the 10-frame short trace. Red spots are the locations of the particle at each frame.

#### VI. DIFFUSION

The MSD curves were fitted by a simple run-and-tumble model $M(\Delta t) = 2v^2\tau\left[\Delta t - \tau(1 - e^{-\Delta t/\tau})\right]$ for evaluating $D^{\rm eff} = v^2\tau/2$ [4]. Figure S10 shows $v$ and $\tau$ obtained by the fitting.

The fitting functions in Fig. 4b are $y(x) = 3800e^{-x^2/230} + 250$ for $h = 5\,\mu$m, $y(x) = 1300e^{-(x-36)^2/38} + 80 + 1600e^{-(x+10)^2/1600}$ for $h = 10\,\mu$m, and $1000e^{-(x-41)^2/200} + 300 - x$ for $h = 20\,\mu$m. Here, $x$ and $y$ are in the unit of cells/10000pix$^2$ and pix$^2$, respectively (1 pix = 0.551 μm).

#### VII. SIMULATION

##### A. Model

We consider a particle-based model where a bacterial cell is expressed by a disk particle with a radius $a$. The particles are self-propelled with a speed $v_0$ along the direction of their polarity $\vec{p}$. The position $\vec{r}_i$ of the $i$th particle and its orientation $\vec{p}_i(t) = (\cos\theta_i(t), \sin\theta_i(t))$ are described by the following dynamical equations:

$$\gamma\left(\dot{\vec{r}}_i - v_0\vec{p}\right) = \sum_{j\neq i}\vec{f}_{ij} \tag{S1}$$

$$\dot{\theta}_i = g_a + g_h - s(\vec{r})\sin(\theta_i - \theta_{\rm laser}) + \xi_i, \tag{S2}$$

where $\gamma$ is the frictional coefficient, and the white noise $\xi_i$ satisfies the statistics of

$$\langle\xi_i(t)\rangle = 0 \tag{S3}$$

$$\langle\xi_i(t)\cdot\xi_j(t')\rangle = 2D_r\delta(t-t')\delta_{ij}. \tag{S4}$$

The variance of the noise is $2D_r$, and expressed by the rotational diffusion constant $D_r$. The rotational Peclet number is given by

$$\mathrm{Pe} = \frac{v_0\tau_r}{a} \tag{S5}$$

and the rotational diffusion time is $\tau_r = 1/D_r$. $a$ is the particle radius. In this study, we set to be $\gamma = 1$, $Dr = 1/6$, and $v_0 = 20/3\sqrt{3/(4\pi)} \approx 3.26$. The Peclet number is Pe $\approx 20$ in accordance with the experimental results. We used $N = 8192$ particles, and varied the system size for each volume fraction.

The first term in (S2) describes the alignment of neighbouring particles within the particle in the region of $\mathcal{B}_i$. We use the polar-type alignment interaction described as

$$g_a = -\sum_{j\in\mathcal{B}_i} g_1\sin(\theta_i - \theta_j) \tag{S6}$$

We choose the interaction range as $r_{ij} < 3a$ and $g_1 = 1$. This term is given by defining the potential energy as

$$U_{\rm align} = -g_1\sum_{r_{ij}<3a}\vec{p}_i\cdot\vec{p}_j. \tag{S7}$$

The distance between the two particles indexed by $i$ and $j$ is denoted by $r_{ij} = |\vec{r}_j - \vec{r}_i|$.

Without the term $g_h$, the system described by (S2) exhibits homogeneous disordered state, motility-induced phase separation (MIPS), and macroscopic polar-order clusters[5]. The MIPS occurs at the high Peclet number, and the polar order appears at stronger alignment. At a lower density, the system shows the disordered state. In our experiments, we did not observe macroscopic density inhomogeneity and macroscopic polar order. There are a number of reasons to their suppression. We include the long-range rotational hydrodynamic interaction between two pushers[6, 7] as

$$g_h = -\sum_{i\neq j}\beta_2\frac{3a^2}{r_{ij}^3}\sin 2\left(\theta_j - \psi_{ij}\right) \tag{S8}$$

where $\psi_{ij} = \arg(\vec{r}_j - \vec{r}_i)$ is a relative angle of the centerline of particles $i$ and $j$. We choose as $\beta_2 = 2$. Because this approximated hydrodynamic interaction is far-field in nature, we take it into account only for the particles with a distance $r_{ij} \geq 3a$. Two closed particles are dominated by the alignment interaction.

We apply the external field $\vec{s}$ expressed by the potential energy $U_{\rm ext} = -\vec{s}\cdot\vec{p}$ leading to the third term in (S2). The excluded volume interactions of translational motion, $\vec{f}_{ij} = -\nabla_{\vec{x}_i}U$ in (S1), arises from the steric repulsive potential, $U$. We use truncated Lennard-Jones to express the repulsion

$$U = \begin{cases}\epsilon\sum_{i\neq j}\left[-2\left(\frac{2a}{r_{ij}}\right)^6 + \left(\frac{2a}{r_{ij}}\right)^{12}\right], & \text{for } r_{ij} < 2a,\\ 0, & \text{otherwise.}\end{cases} \tag{S9}$$

The steric interaction avoids divergence due to the hydrodynamic interaction when two particles overlap. In this study, we consider both with the steric interaction $\epsilon = 1$ and without it $\epsilon = 0$. The latter imitates the system with a larger height in which particles can pass each other.

##### B. Response to an external field

We consider responses of particle motions under an external field. In this study, we consider two types of external fields; one is a uniform external field under which particles are aligned in the direction of $\theta_0$ (Fig.6b). The other one is a Gaussian-shaped external field, under which each particle tends to align along the direction of $\theta_{\rm laser} = \arg\vec{r}/|\vec{r}|$ (Fig.6a).

Under the uniform external field, we measure the probability distribution of the orientation of each particle, as shown in Fig.6b in the main text. Because the external field forces particles pointing toward $\theta = 0$, the distribution has a peak there. At a small volume fraction of particles, the peak is broader, suggesting that particles do not respond to the uniform external field. On the other hand, at a large volume fraction, the particles respond strongly to the external field.

We also consider the Gaussian-shaped external field to model the experimental setup. Similar to the experiments, we apply the external field in the form of the Gaussian function, $s(\vec{r}) = s_0 e^{-|\vec{r}^2|/2w^2}$ with the width of $w = 100a$. The particles are aligned along the radial direction of the external field, namely $\theta_{\rm laser} = \arg \vec{r}/|\vec{r}|$ with its strength $s(\vec{r})$. We choose the amplitude of the external field to be $s_0 = 0.1$. After switching on the external field, the particles start to accumulate near the origin because they tend to point toward the center of the external field. We measure the density $\rho(t)$ inside the region of $|\vec{r}| < w$. The response to the external field is computed by the gradient of the $\rho(t)$ during the time window $\Delta t \approx 166\tau_r$ after switching on the field. The result is shown in Fig.6a of the main text. Here, we use the simulation data of 3-20 independent runs.

*Mean-field analysis of the response to an external field*

In this section, we discuss the origin of the density-dependent response using a mean field approximation. We consider an idealized system where the external field is applied uniformly in space. We also neglect the effect of the long-range rotational interaction $g_h$. Under the mean-field approximation, all the particles are on average aligned in the direction of the external field, and thus, (S2) is written as

$$\dot{\theta}_i = -ng \sin(\theta_i - \theta_{\rm laser}) - s \sin(\theta_i - \theta_{\rm laser}) + \xi_i. \quad \text{(S10)}$$

Here, $n$ is the number of interacting particles. We choose the external field along the $x$ axis $\vec{e}_x$ without loss of generality, and thus $\vec{s} = (s, 0)$ and $\theta_{\rm laser} = 0$. With this simplified model, the probability distribution of the orientation is given by

$$P(\theta_i) = Z^{-1} e^{-\beta H} \quad \text{(S11)}$$

$$Z = \int e^{-\beta H} d\theta_i \quad \text{(S12)}$$

with the effective free energy

$$H = -(ng + s) \cos\theta_i \quad \text{(S13)}$$

under the inverse temperature $\beta = \frac{1}{D_r}$.

In Fig. 6b, we compare the orientation distribution of particles in the simulations and that is evaluated from (S11). The orientation is identified by the argument of velocity vector $\tan^{-1} v_y/v_x$. Both simulation data and theoretical results show the distribution of the orientation is accumulated at $\theta = \theta_0$ close to the direction of the external field at higher density. When the density is low, the mean-field approximation fits the simulation data. At the higher density, the mean-field approximation qualitatively reproduce the distribution, but it overestimates. This is due to correlation effects between particles.

##### C. Effective diffusion

We consider the effective diffusion constant without an external field, $s = 0$. We measure the mean-square displacement (MSD)

$$M(\Delta t) = \langle |\vec{r}_i(t + \Delta t) - \vec{r}_i(t)|^2 \rangle, \quad \text{(S14)}$$

where the average is taken over all the particles and time at the steady state. The MSD was fitted by Eq. (2) to obtain $D^{\rm eff}$.

##### D. Effective Soret coefficient

The effective Soret coefficient $S_{\rm T}^{\rm eff}$ increased with $\varphi$ (Fig. S5), which qualitatively reproduced the experimental result (Fig. 5). However, the calculation of response and effective diffusion was limited to relatively small $\varphi$ (Fig. 6). Hence, the estimation of $S_{\rm T}^{\rm eff}$ at large $\phi$ may not be reliable.

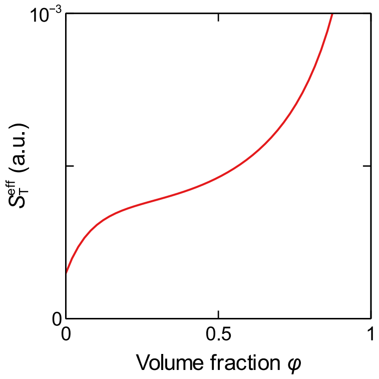


FIG. S5: Effectie Soret coefficient calculated as the ratio of the fitting curves of the response (Fig. 6a) and $D^{\rm eff}$ (Fig. 6c).

#### VIII. SUPPLEMENTARY FIGURES

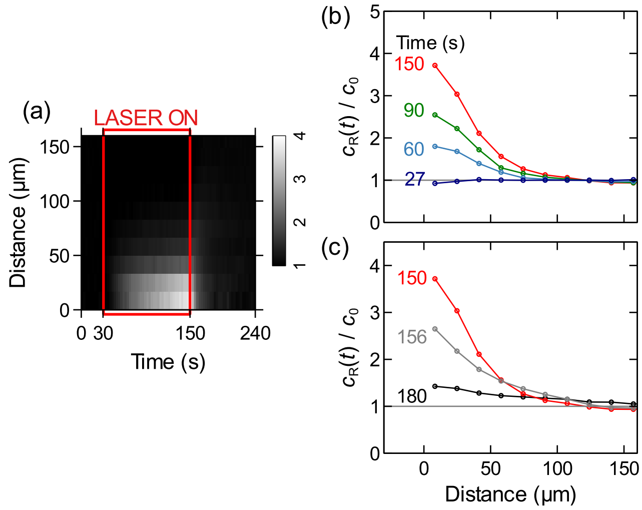


FIG. S6: Spatial profile of cell density. (a), Spatio-temporal distribution of the normalized cell density $c_R(t)/c_0$. (b, c), The Spatial profile at different timing (laser is on during 30 and 150 s).

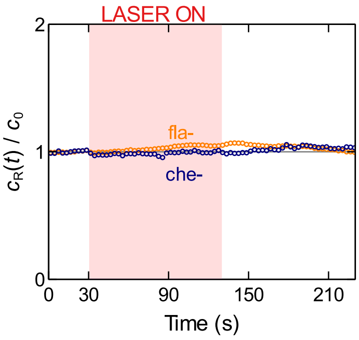


FIG. S7: Taxis of the immotile strain lacking flagella (fla-, 4.1 cells/100μm$^2$) and the nontactic strain lacking Che system necessary for taxis (che-, 3.4 cells/100μm$^2$). The time course of the normalized cell density $c_R(t)/c_0$ in the region within $R$ = 50 μm indicated as a yellow dotted circle in Fig. 2(a). The immotile fla- cells responded to the laser spot slightly, possibly due to the weak optical tweezers effect. che- is motile and not trapped by the optical trapping.

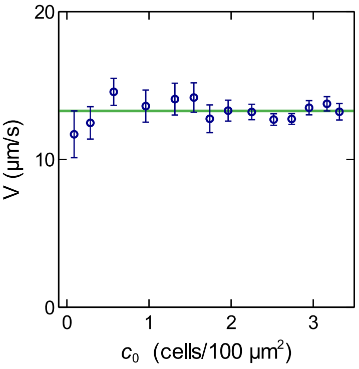


FIG. S8: Mean swimming velocity corresponding to Fig. 2(d, e). There was no significant correlation between $c_0$ and the mean velocity. The error bars correspond to standard errors.

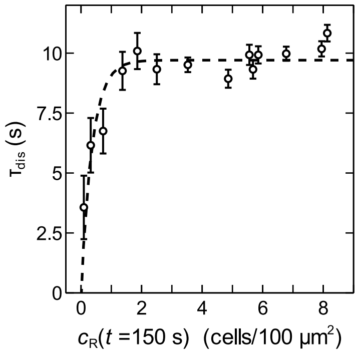


FIG. S9: Time constant of the dispersion process after the laser is turned off is plotted against the cell density at the moment of the laser off ($t$ = 150 s). We fitted $c_R(t)/c_0$ by an exponential function $f(t) = a + b\exp(-\frac{t-150\,\mathrm{s}}{\tau_{\mathrm{dis}}})$ with the fitting parameters $a$, $b$, and $\tau_{\mathrm{dis}}$ in $t \geq 160$ s. The error bars indicate the fitting errors and correspond to standard errors.

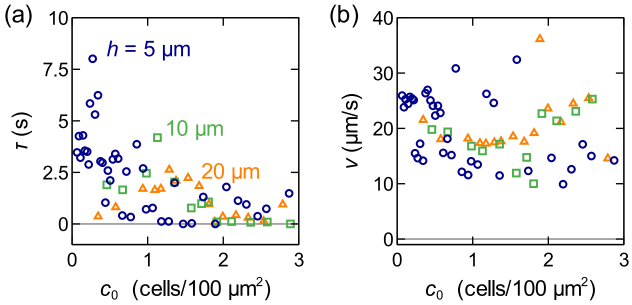


FIG. S10: The values of the fitting parameters $v$ and $\tau$ of the diffusion experiments.

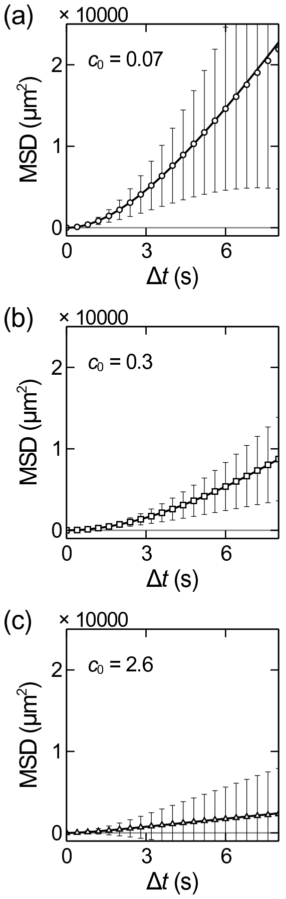


FIG. S11: MSD curves in Fig. 4b with error bars indicating standard deviations.
